# Validation of Dynamic Bayesian Optimization for Human-in-the-Loop Optimization of Exoskeleton Control at User-Driven Walking Speed

**DOI:** 10.64898/2026.06.10.731447

**Authors:** GilHwan Kim, Fabrizio Sergi

## Abstract

Human-in-the-loop optimization (HILO) is an established method for identifying subject-specific optimal controllers for performance augmentation. For HILO algorithms to be useful in rehabilitation, however, the optimization algorithm may need to account for how the human response changes over time in response to assistance.

In this study, we tested a modified version of Bayesian optimization (BO), dynamic Bayesian optimization (DBO), in a three-parameter optimization problem that sought to identify participant-specific optimal solutions for increasing walking speed. As opposed to BO, DBO accounts for the non-stationarity of human responses. Sixteen healthy participants received bilateral hip torque pulses delivered by a hip exoskeleton. The exoskeleton torque parameters were determined using HILO with either DBO or BO. Validation iterations were introduced to objectively compare performance across optimizers at different time points of HILO.

The results showed that both DBO and BO significantly increased walking speed compared to baseline. When comparing performance between DBO and BO, DBO emerged as an improvement over BO both in terms of efficacy, modeling accuracy, and personalization. DBO induced changes in walking speed relative to baseline that exceeded those induced by BO at three of the four validation iterations. DBO outperformed BO in modeling accuracy in later validation iterations. DBO personalization induced changes in walking speed that were significantly greater than those induced by previously identified assistive solutions, while this was not the case of BO. Overall, our results indicate that DBO outperformed BO due to its greater ability to account for non-stationary aspects of the human response.

## I. INTRODUCTION

Robot-assisted gait training, with its ability to deliver consistent and repeatable mechanical assistance, has emerged as a promising approach for rehabilitation following neurological injury [1], [2]. When an appropriate level/type of robotic intervention is applied, desired training effects can be induced. However, due to inter-individual variability, solutions derived from group-averaged responses may result in suboptimal outcomes or no measurable improvement for a given individual. Therefore, effective training requires the construction of personalized input–output relationships to accurately estimate participant-specific solutions that induce the desired training effect. This approach, however, necessitates multiple observations under different control input parameters, which introduces the risk of fatigue, particularly when the number of parameters is large.

To address participant variability and efficiently identify optimal solutions, human-in-the-loop optimization (HILO) has been introduced in previous studies targeting primarily human performance augmentation [3], [4], while using various optimization algorithms [5], [6], [7]. In HILO, the optimizer iteratively updates its estimate of the system response as new observations are collected, enabling efficient identification of individualized response models and optimal control parameters in real time. However, most previously used optimizers assume that the human–robot system is stationary and does not change over time, due to the history of exposure to robot intervention. Under this assumption, the same control inputs applied by the training device are expected to produce similar outputs regardless of when they are applied. This limitation can degrade the accuracy of system estimation and lead to suboptimal or incorrect input selection in cases where the system is highly non-stationary [8].

In rehabilitation, instead, the main goal is to expose participants to training, aiming to induce *changes* in motor coordination that may be beneficial to walking function if they transfer beyond training. For the success of HILO algorithms to rehabilitation, it is essential to adopt optimization algorithms capable of handling the non-stationary nature of the human–robot system. Furthermore, prior HILO studies in robot-assisted gait training have primarily focused on minimizing metabolic cost, which has limited direct relevance to rehabilitation goals. More direct biomechanical outcomes, such as self-selected walking speed or metrics of propulsion mechanics, may provide more meaningful targets for optimization.

Dynamic Bayesian optimization (DBO), an extension of conventional Bayesian optimization (BO), has been proposed to address non-stationarity in optimization problems. DBO operates by modifying the conventional BO algorithm by adjusting the Gaussian Process model used for estimating the system’s response to account for the time at which different observations where collected. In general, for HILO, we can assume that, compared to recent observations, older observations are equally or less predictive of the future system response due to changes in the system properties that may have occurred with exposure to robot assistance. To account for the unknown time-varying capabilities of the system’s response, a smooth decay function has been introduced when constructing the Gaussian process model [9]. Several studies showed that dynamic Bayesian optimization outperforms existing optimization methods including conventional Bayesian optimization [10], [11], [12], [9], [13], [14] in finding optimal solutions for systems with explicitly time-varying input-output relationships. Recent work has explored the application of DBO within HILO settings, including simulations of neuromotor learning [8], pilot experimental studies in robot-assisted gait training [15], and robot-assisted gait training targeting propulsion mechanics [16]. These studies reported performance similar to or exceeding that of standard BO. However, these implementations have been limited to numerical simulations or to experimental conditions with artificially introduced non-stationary aspects in the human response, intended primarily for proof of feasibility. As a result, DBO has not yet been fully evaluated in practical HILO-based robot-assisted gait training for achieving biomechanically meaningful training outcomes, such as improvements in walking speed or propulsion.

In this study, we implemented DBO within a multi-parameter human-in-the-loop robot-assisted gait training framework targeting increase of walking speed. We conducted a human-subject experiment to determine whether DBO could induce changes in participant walking speed during training, and whether the changes induced by DBO were greater than those achievable with BO or with a sham condition where robotic inputs were delivered with the same variability experienced during the DBO and BO sessions. Sixteen healthy participants were exposed to the action of a bilateral hip exoskeleton applying torque pulses to participants walking on a treadmill under user-driven control, where belt speed was adjusted according to the participant intention and action. The torque profiles delivered by the hip exoskeleton were selected via dynamic Bayesian optimization or conventional Bayesian optimization during real-time HILO towards the goal of maximizing walking speed within reasonable safety constraints.

## II. Methods

### A. Dynamic Bayesian Optimization

Dynamic Bayesian optimization (DBO) is a version of Bayesian optimization (BO), developed to account for the dynamic nature of an objective function [17], [8], [9]. A detailed explanation is provided in [8], but a brief overview is presented in this section to facilitate understanding of the experimental work described in this paper.

In conventional BO, a Gaussian process model is constructed at each iteration to make a prediction of the system output. In our case, the system is the human-robot system, and the output is a metric of biomechanical performance resulting from human-robot interaction. Based on this model, BO selects the input value to be evaluated in the next iteration. The Gaussian process model can be described as:

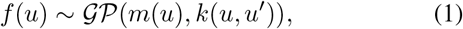

where *m*(*u*) = 𝔼 [*f* (*u*)] is the mean function, representing the expected value of the objective function at input *u*, and *k*(*u, u*′) is a covariance function, which characterizes the relationship between objective function values at two different input points. Since conventional BO assumes that the objective function does not change over time or across iterations, the Gaussian process model inevitably exhibits limited prediction accuracy when this algorithm is applied to systems with a dynamic component. This limitation becomes more pronounced when human factors, such as neuromotor learning and adaptation behaviors, are involved [18].

To account for the dynamic aspects of the system, DBO is introduced by incorporating time as an additional variable *t* in the covariance function, i.e., *k*((*u, t*), (*u*′, *t*′)), where *t* and *t*′ denotes the time instances corresponding to the inputs *u* and *u*′, respectively. To simplify this covariance function, prior implementations of DBO assumed separability between the dynamic and the static component of the covariance function [9], [19]. Under this assumption, the overall covariance function *k*((*u, t*), (*u*′, *t*′)) can be expressed as the product of a static *k*_*u*_(*u, u*′) and dynamic component *k*_*t*_(*t, t*′):

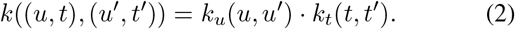

Based on the rationale that older observations may mislead the estimation of the true objective function [9], the dynamic component of covariance function, *k*_*t*_(*t, t*′), is defined as smooth decaying function related to the time difference as:

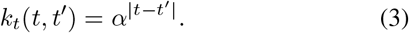

where the hyperparameter *α* ∈ (0, 1] defines the time scale, determining the covariance between measurements collected at times *t* and *t*′ (Fig. 1b). In this work, the static component of the covariance function, *k*_*u*_(*u, u*′), was implemented as the automatic relevance determination squared exponential function [20].

**Fig. 1.**
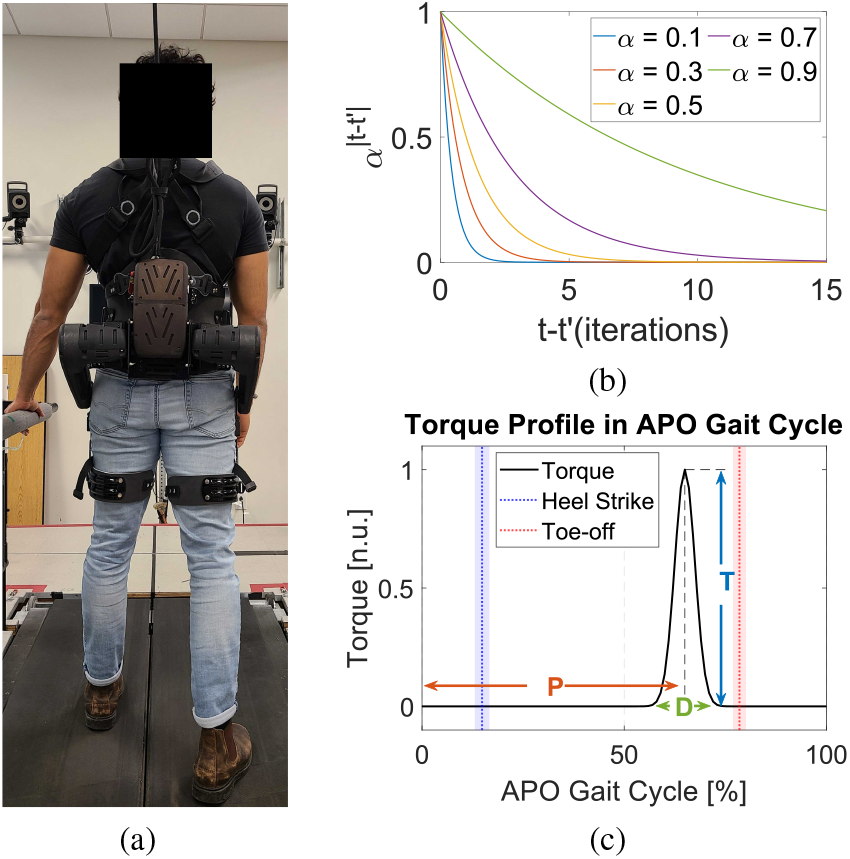
(a) Experimental setup, including the active pelvis orthosis (APO), instrumented treadmill, motion capture system, and safety harness. (b) Dynamic component of the covariance function, with each line representing the covariance function with different hyperparameter *α*. (c) Hip torque profile in the APO Gait Cycle domain, where 0% corresponds to peak of hip flexion. Heel-strike and toe-off timings are shown as distributions, with the mean and standard deviation indicated in blue and red, respectively. The control parameters defining the torque profile (duration: *D*, timing: *P*, and amplitude: *T*) are labeled.

#### 1) Periodic control parameter

In this work, we extend the optimization framework to include multiple control parameters and to account for the periodicity of some control parameters. The parameters are illustrated in Fig. 1c. *P* is a periodic input parameter as it encodes pulse application timing in the normalized 0-100 domain of percent gait cycle. As such, the metric of distance between two input values that is intrinsic to the variance function needs to be modified to account for the periodicity of control parameters. In our work, we modified the covariance function as

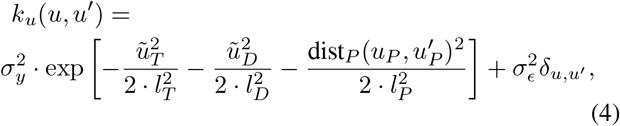

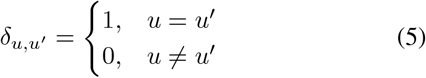

where 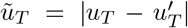 and 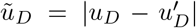 are abso-lute difference in input value of *T* and *D* respectively. The distance metric for the periodic parameter *P* is defined as dist_*P*_ (*u, u*′) = min(|*u − u*′ |, 100 − |*u − u*′|), which represents the minimum distance between two values of the periodic input parameter with periodicity equal to 100 (units of % gait cycle duration). The hyperparameters *l*_*T*_, *l*_*D*_, and *l*_*P*_ correspond to the length scales associated with *T, D*, and *P*, respectively. 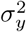 denotes the variance of the measurements, while 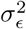 represents variance due to noise. Therefore, the overall covariance function is denoted as below.

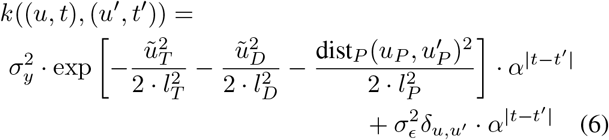

### B. Experimental Methods

#### 1) Equipment

The Active Pelvis Orthosis (APO), a bilateral lower extremity robotic exoskeleton, was used to apply torque pulses to participants during the experiment (Fig. 1a). The APO (IUVO, Pisa, Italy) can generate hip torque pulses to both hip joints [21], with a peak torque of 10 Nm, and is equipped with onboard encoders that measure the hip joint angles with a resolution of 0.015 deg and a sampling rate of 100 Hz. Based on the hip joint angle measured in the previous gait cycle, with 0 % defined as the point of maximum hip flexion, the APO estimates the participant’s current gait phase using adaptive oscillators [22]. The gait phase is estimated separately for the left and right legs, and is used to define the timing of hip torque pulses in Gaussian function shaped profile, defined by three control parameters: torque pulse amplitude (*T*), duration (*D*), and peak timing (*P*) (Fig. 1c).

In this experimental study, the control parameters (*T, D*, and *P*) were updated iteratively in real-time via Simulink (MathWorks Inc., Natick, MA, USA). Via a Serial Peripheral Interface connection, the Simulink controller sends updated control parameters to the APO and receives measurements of hip joint angle and torque pulse magnitude. To ensure proper execution of the updated torque profile, the new setting of control parameters to be used in the next stride were updated and transmitted to the APO during the wash-out phase of previous iteration, where no torque was applied to the participant.

A split-belt instrumented treadmill (Bertec Corp., Columbus OH, USA) was used in this experiment with two force plates integrated into the treadmill. Force plate data from both treadmill belts were sampled in real-time at a frequency of 400 Hz and were used to adjust treadmill speed through a user-driven treadmill controller, in which treadmill velocity changed in response to participant walking behavior [23]. For safety, the maximum treadmill speed was limited to 1.6 m/s, and a harness system (Solo-Step Inc., North Sioux City, SD, USA) connected to an overhead railing track was used to prevent the participant from falling.

#### 2) Study Participants

Sixteen healthy participants (9 females; age (mean ± std): 24.4 ± 3.4 yrs, height: 170.3 ± 8.5 cm, and mass: 70.1 ± 9.2 kg) were recruited for this experiment (protocol no. 1755609-9, approved by the University of Delaware Institutional Review Board). Each participant completed two experiments on different days, with a minimum interval of one day and a maximum interval of seven days between sessions. The two experiments employed different optimization algorithms (DBO vs. BO), and the order in which the optimizers were assigned to participants was pseudorandomized and evenly balanced across the participants.

#### 3) Training Procedures

During each experimental visit, the protocol consisted of a preparation session and of a familiarization session, followed by the main training session. During the preparation session, the APO was fitted to the participant, ensuring proper alignment between the participant’s hip joint axis and the APO’s rotational axis in sagittal plane. After confirming the participant’s comfort, participant was secured with the safety harness.

During the familiarization session, the participant first walked on the treadmill under user-driven treadmill control to find their comfortable walking speed, with the robot in zero-torque mode. After at least 3 minutes of walking, once a stable walking speed was observed, the participant was exposed to three different torque pulses applied to both hip joints while walking on the treadmill. Torque pulses were generated based on combinations of three control parameters: torque amplitude (*T*), duration (*D*), and timing (*P*). In the familiarization session, three sets of torque pulses were used: two assistive hip torque pulses targeting the stance and swing phases of walking, with parameter sets (*T, P, D*) = (−9, 75, 30) and (9, 50, 40), respectively, and one resistive torque pulse applied during the stance phase, with parameters (*T, P, D*) = (−5, 50, 20). The assistive torque pulse parameters were selected based on findings from a previous pilot study conducted using the same experimental setup [15]. During the familiarization session, the order of the torque pulses was randomized across participants. Across training, each new torque pulse candidate was applied gradually, starting with a duration (*D*) of 10%. The torque amplitude was then increased in steps of 2 Nm every two strides until reaching a maximum amplitude of 9 Nm for each parameter combination. After reaching this maximum amplitude, the pulse duration was gradually increased to the desired value, in increments of 15% every two strides, until the value targeted by HILO was applied. The gradual increase of torque amplitude and duration took place over a maximum of 14 strides in all participants. During the second experimental visit, participants completed the same familiarization session as in the first visit.

During the main experiment session, participants were exposed to four training trials, each lasting up to 15 minutes, with a minimum 5-minute rest period between trials. In each training trial, participant experienced 10 different torque pulse conditions to both hip joints while walking on the treadmill, which was controlled via a user-driven treadmill controller. In the first trial, the first three torque pulses were predetermined and consistent across participants (see details in Sec. II-B4). The torque pulses were introduced gradually, with a maximum increase of 2 Nm per stride. Each torque pulse condition was applied continuously for 40 strides, followed by a 20-stride wash-out phase during which no torque was applied to the participant. In the first trial of training, the participant began with a 100-stride baseline phase with no torque was applied, used to identify the preferred walking speed. After each training trial, the harness was removed, and participants were asked to sit and rest for at least 5 minutes, ensuring they were free of fatigue before beginning next trial. The subsequent trial began at the final speed of the previous trial, with the user-driven controller activated after 10 strides to maintain that speed and adjust according to user input.

An additional experimental visit was conducted for three participants under a sham torque condition. This visit followed the exact same procedure as the previous experiments that used either the DBO or BO optimizer. The key difference was in the torque applied during the main training session: from the 4^th^ to the 40^th^ iteration, the participant experienced a random torque condition. These torque values were generated from a distribution with a randomly selected mean of the input torque parameters (*T, P, D*), while maintaining the same covariance observed in the sessions conducted with DBO or BO. This procedure ensured that the applied torque had random mean value, but had similar variability in this sham condition as those identified by BO and DBO in the main experiment. Specifically, a covariance matrix was derived from the input parameters spanning the 4^th^ to 40^th^ iterations of the main experiment results using DBO or BO as optimizer, and the matrix with the higher variance was selected to define input parameters for sham condition. Using this covariance structure and a randomly sampled torque pulse parameter vector, 37 additional input parameter sets were generated to preserve the same distribution. These input sets were then used as input for the 4^th^ to 40^th^ iterations of sham condition, while the 1^st^ to 3^rd^ iterations used the same input parameter values as in the main experiments.

#### 4) Optimizer Setup

The optimization methods, DBO or BO, were implemented during the training sessions to determine the control parameters for the hip torque profile at each iteration. The objective of both DBO and BO was to maximize the participant’s walking speed by minimizing the cost function *v*, where *v* represents the treadmill speed. In the Results section, walking speed*− v* is reported for clarity and ease of interpretation. The average walking speed over the last five strides of the torque intervention phase at each iteration (strides 36–40) was used as the measured outcome for that iteration. During each iteration, the participant experienced the same torque pulse input, except for the first few strides, during which the torque amplitude was gradually increased, and the final 20 strides of the washout phase, when no torque was applied.

The range of torque amplitude (*T*) was from -9 Nm (maximum flexion) to 9 Nm (maximum extension), peak timing (*P*) ranged from 0 % to 100 % of the APO gait cycle, and duration (*D*) ranged from 10 % to 40 % of gait cycle. The initial three inputs used in the first training trial were predefined as (*T, P, D*) = (−9, 75, 30), (9, 50, 40), and (−5, 50, 20), representing assumed solutions for swing and stance assistance, as well as one resistance condition applied during the stance phase of the gait cycle. Expected improvement was used as acquisition function, as it demonstrated the highest performance in pervious simulation study [24]. The exploration and exploitation ratio was set to 0.1, which favoring exploitation, based on a previous simulation [8] and pilot test result [15], to reduce excessive variability and provide a more intuitive experience for the participant.

A total number of 40 iterations, (i.e., 10 iterations per each training trial), were performed in each main experiment session. A validation iteration was conducted at every 10^th^ iteration (10^th^, 20^th^, 30^th^, and 40^th^ iteration), which was the final iteration of each training trial, during which the optimizer tested its estimated best input parameters to maximize walking speed (min predicted mean value of the estimated model). At the beginning of the next training trial, the optimizer used all previous observations, including those from the validation iterations, to continue the optimization process. The prediction error was calculated as the absolute difference between the observation in the validation iteration and the corresponding response predicted by the mean function of GP model, which was constructed using all prior observations:

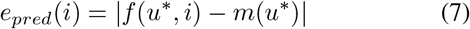

where *e*_*pred*_(*i*) denotes the prediction error at iteration *i. u*\* is the input parameter set evaluated at iteration *i*. The function *f* (*·,·*) represents the measured output of the human–robot system, while *m*(*·*) denotes the mean function of the Gaussian Process model constructed using observations from iterations 1 to *i −*1. The standardized prediction error was also calculated based on the prediction error as:

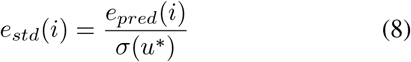

where *σ*(*u*\*) is the standard deviation (uncertainty) of the GP model at input *u*\*.

#### 5) Statistical Analysis

As participants were exposed to two different optimizer conditions and their response evaluated longitudinally across each experiment, the responses at each validation iteration were collected and compared across different optimizer condition. Linear mixed-model analyses were conducted with optimizer type, iteration, and their interaction included as fixed effects. Random effects included participant and the interaction of participant with optimizer type and iteration. Walking speed and prediction error were analyzed separately using linear mixed models to quantify effectiveness of the assistance of BO and DBO, and their modeling accuracy, respectively. For walking speed, iteration was modeled with five levels (baseline and validation iterations), whereas for prediction error, iteration included four levels (validation iterations only). When significant main effects or interactions were observed, post hoc pairwise t-tests were performed to compare all combinations of optimizer type and iteration number. Effect sizes were reported as standardized mean differences (Cohen’s *d*_*z*_ for paired t-tests and Cohen’s *d* for comparisons against a constant).

## III. Results

### A. Walking speed

The linear mixed model analysis was conducted to analyze longitudinal changes in walking speed to evaluate the effects of optimizer type (DBO vs. BO) across iterations. The model results on the walking speed outcome revealed a significant main effect of iteration number (*p <* 0.0001) and of the interaction between optimizer type and iteration (*p* = 0.015). In contrast, optimizer type (*p* = 0.150) did not have a significant effect on walking speed.

#### 1) Optimizer-induced increase in walking speed

Walking speed data were compared by each optimizer type at multiple iterations (Fig. 2). Both optimizers showed an increase in speed at the final iteration, relative to baseline. Paired comparisons revealed that walking speed was significantly different from baseline at iterations 1, 2, 30, and 40, consistently across optimizer types. For DBO, the changes were 0.08 ± 0.03 m/s (*p* = 0.03, *d*_*z*_ = 0.615), 0.11 ± 0.03 m/s (*p* = 0.005, *d*_*z*_ = 0.841), 0.30 ± 0.04 m/s (*p <*.0.0001, *d*_*z*_ = 1.94), and 0.30 ± 0.04 m/s (*p <* 0.0001, *d*_*z*_ = 1.90) at iterations 1, 2, 30, and 40, respectively. For BO, the corresponding changes were 0.08 ± 0.03 m/s (*p* = 0.012, *d*_*z*_ = 0.741), 0.11 ± 0.03 m/s (*p* = 0.001, *d*_*z*_ = 1.011), 0.15 ± 0.03 m/s (*p <*0.001, *d*_*z*_ = 1.124), and 0.18 ± 0.06 m/s (*p* = 0.007, *d*_*z*_ = 0.803), respectively. Towards quantifying the benefit of participant-specific optimization, we sought to compare the effects of the assistance candidates applied in iterations 1 and 2 (defined empirically based on pilot testing), with the candidates tested in cross-validation iterations 30 and 40, i.e., after approximately 1800-2400 strides of exposure to HILO. As shown in Fig. 2, DBO resulted in a significant increase at iteration 30 and 40 compared to iteration 1 (iteration 30: 0.23 ± 0.05 m/s, *p <* 0.001, *d*_*z*_ = 1.171; iteration 40: 0.22 ± 0.05 m/s, *p <* 0.001, *d*_*z*_ = 1.161), as well as compared to iteration 2 (iteration 30: 0.20 ± 0.04 m/s, *p <* 0.001, *d*_*z*_ = 1.249; iteration 40: 0.19 ± 0.05 m/s, *p* = 0.001, *d*_*z*_ = 1.014). In contrast, BO did not induce a significant difference in walking speed between late validation iterations and iteration 1 (iteration 30: 0.07 ± 0.03 m/s, *p* = 0.06; iteration 40: 0.11 ± 0.06 m/s, *p* = 0.09), or iteration 2 (iteration 30: 0.04 ± 0.04 m/s, *p* = 0.37; iteration 40: 0.08 ± 0.06 m/s, *p* = 0.24).

**Fig. 2.**
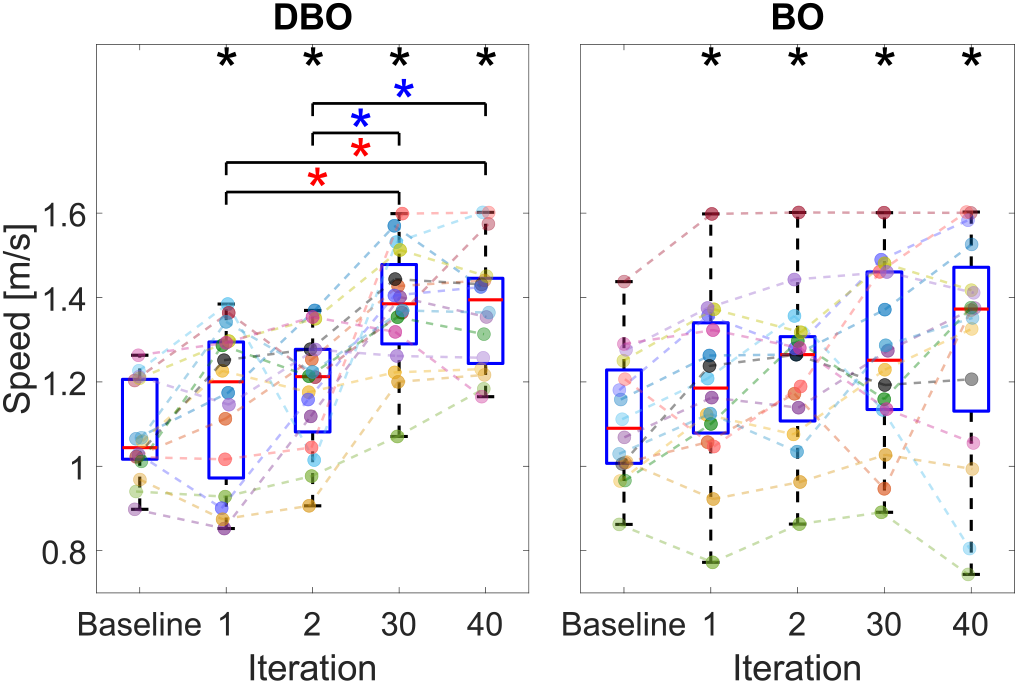
Walking speed resulting from either optimizer across iterations. Paired comparisons were conducted separately for each optimizer type between iterations. Black asterisks indicate significant differences relative to baseline (*p <* 0.05). Red and blue asterisks indicate significant differences relative to iteration 1 and 2, respectively. In iterations 1 and 2, assistive torques assumed to increase walking speed via swing or stance phase assistance, respectively, were applied.

#### 2) Performance difference between DBO and BO

Group averaged walking speed data at each iteration are shown in Fig. 3, with individual participant responses in Fig. 4. To facilitate between-optimizer comparisons, group-averaged walking speed changes relative to the specific session baseline are shown in Fig. 3, bottom. Under both DBO and BO, walking speed generally increased over iterations, and it was significant higher than baseline across all validation iterations. In contrast, the mean values for the sham condition remained largely unchanged, staying relatively constant throughout the experiment, and did not differ significantly from baseline across all validation iterations. During each validation session, walking speed under sham torque condition remained approximately constant, with the values of 1.1 ± 0.1 m/s, 1.1 ± 0.1 m/s, 1.2 ± 0.1 m/s, and 1.1 ± 0.1 m/s for iterations 10-40, respectively.

**Fig. 3.**
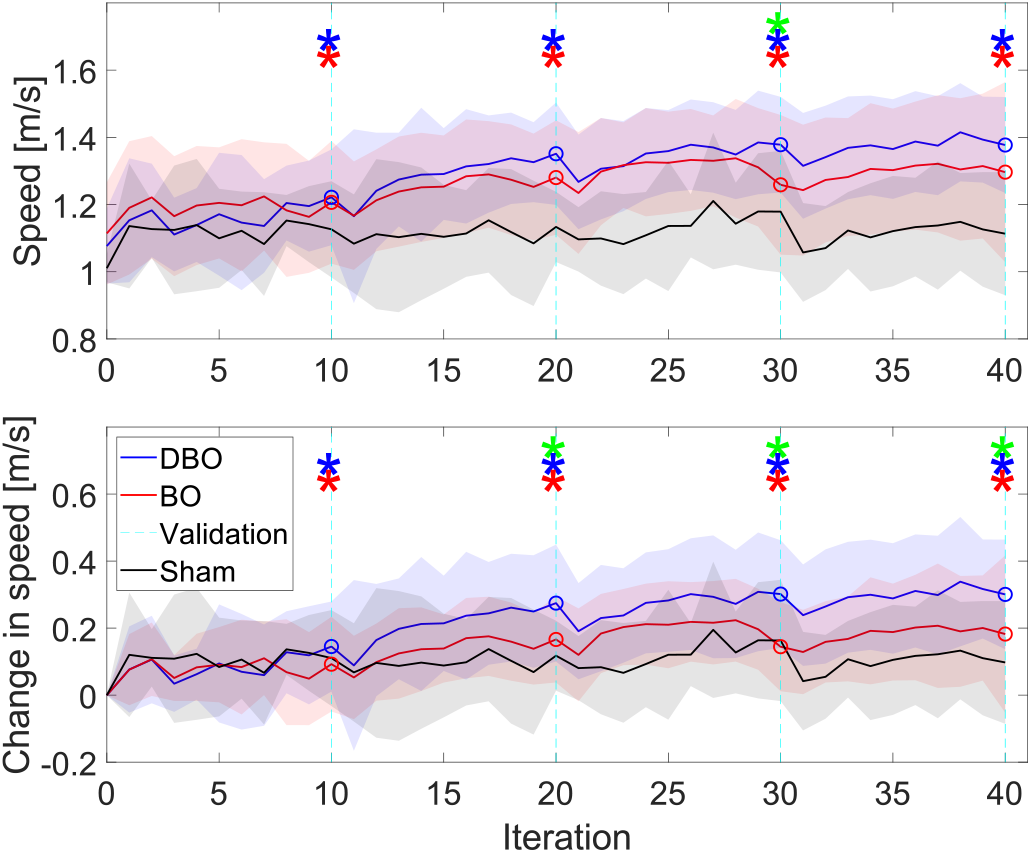
Group-level walking speed across the experiment (top: walking speed, bottom: change in walking speed relative to baseline). The solid lines represent the mean walking speed for 16 participants at each iteration for DBO (blue), BO (red), and for three participants in the sham condition (black). The shaded area indicate 1 ± standard deviation with corresponding colors for each condition. Responses at 0^th^ iteration correspond to observations at baseline. The cyan dashed line indicates validation iterations where the estimated best input was tested, corresponding walking speed function values are shown with a circle marker. A green asterisk denotes significant difference between DBO and BO (*p <* 0.05), while red and blue asterisks indicate significant changes in walking speed relative to baseline and at each iteration.

**Fig. 4.**
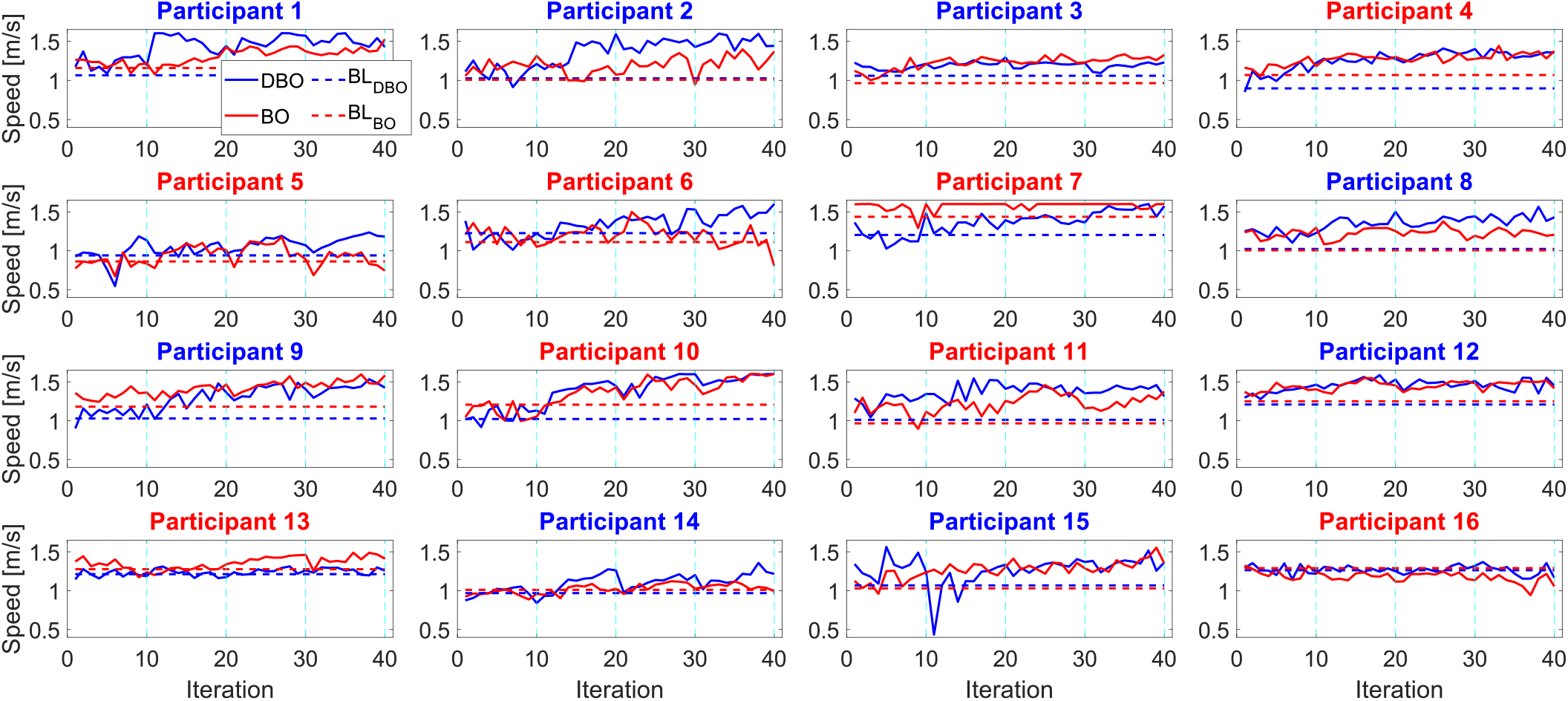
Individual participant walking speed across the experiment. Blue solid lines represent participant responses using DBO, while red lines correspond to BO. The vertical cyan dashed lines indicate the validation iterations. Dashed line extend to visually indicate the baseline speed for each session, color-coded by optimizer type. The color in the panel title text indicates which optimizer was tested first in a given participant.

Changes in walking speed relative to baseline were compared at each validation iteration between optimizer type (DBO and BO). DBO showed significantly greater increases in walking speed at iterations 20 (0.11 ± 0.04 m/s, *p* = 0.013, *d*_*z*_ = 0.724), 30 (0.16 ± 0.04 m/s, *p* = 0.002, *d*_*z*_ = 0.999), and 40 (0.12 ± 0.05 m/s, *p* = 0.04, *d*_*z*_ = 0.579). When looking at the absolute values of walking speed resulting from exposure to the two optimizers (i.e., not the baseline-subtracted change), only the comparison at iteration 30 showed a significant difference in walking speed between DBO and BO (DBO: 1.38 ± 0.04 m/s, BO: 1.26 ± 0.05 m/s, difference: 0.12 ± 0.05 m/s, *p* = 0.02, *d*_*z*_ = 0.648). However, at the final iteration, the difference between DBO and BO was no longer significant (0.08 ± 0.06 m/s, *p* = 0.21).

### B. Model accuracy

Model accuracy was assessed by analyzing of the prediction error at the validation iterations, during which the optimizers applied the assistance candidate predicted to produce the greatest walking speed for each participant. The linear mixed model results on the prediction error revealed a significant main effect of the interaction between optimizer type and iteration (*p* = 0.030). In contrast, optimizer type (*p* = 0.374) not iteration number (*p* = 0.374) had a significant effect on prediction error. Prediction error data were further compared across validation iterations for each optimizer type (Fig. 5). For prediction error, DBO showed a significant reduction at iteration 30 compared to iterations 10 and 20 (iteration 10: 0.06 ± 0.02, *p* = 0.023, *d*_*z*_ = 0.357; iteration 20: 0.03 ± 0.01, *p* = 0.022, *d*_*z*_ = 0.659). In contrast, BO showed no significant differences across validation iterations. Similarly, for standardized prediction error, DBO showed a significant reduction at iteration 40 compared to iterations 10 and 20 (iteration 10: 9.29 ± 3.20, *p* = 0.011, *d*_*z*_ = 0.751; iteration 20: 4.80 ± 2.04, *p* = 0.033, *d*_*z*_ = 0.607), whereas BO again showed no significant differences across validation iterations.

**Fig. 5.**
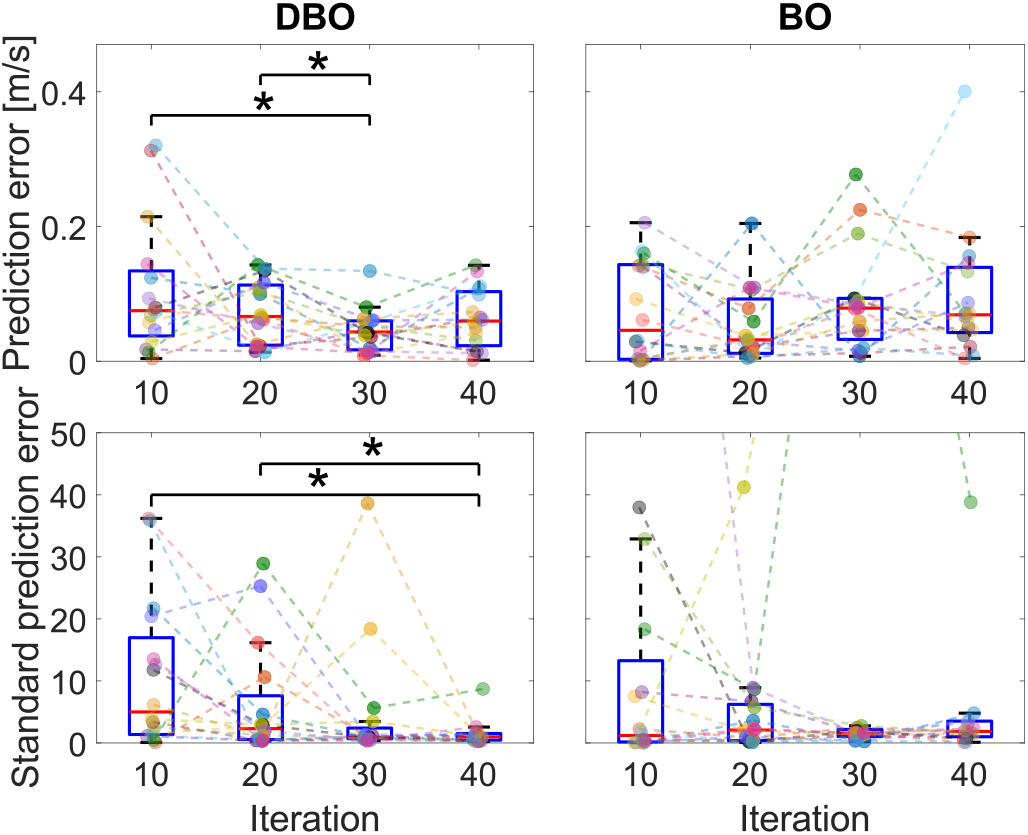
Paired comparisons broken down by iteration number and optimizer type. Colored dots indicate each participant’s data. An asterisk represents a significant difference based on paired comparisons.

As shown in Fig. 6 top, the prediction error for DBO, along with its standard deviation, decreased over iterations, with values of 0.10 ± 0.02 m/s, 0.07 ± 0.01 m/s, 0.05 ± 0.01 m/s, and 0.06 ± 0.01 m/s for iterations 10-40. In contrast, BO maintained a relatively consistent prediction error throughout the experiment, with values of 0.07 ± 0.02 m/s, 0.05 ± 0.01 m/s, 0.09 ± 0.02 m/s, and 0.10 ± 0.02 m/s for the same iterations.

**Fig. 6.**
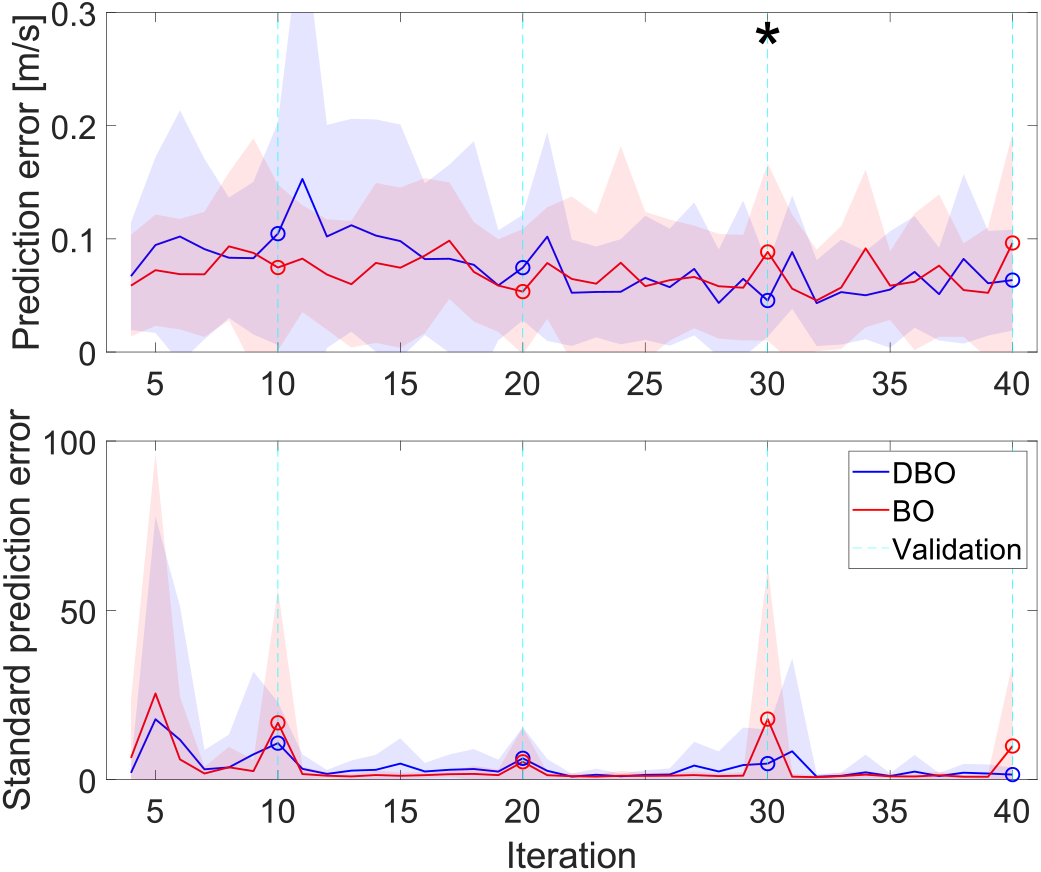
Group-level prediction error across the experiment (top: prediction error, bottom: standard prediction error). The asterisk denotes significant difference between DBO and BO (*p <* 0.05).

Paired comparisons between optimizer conditions (DBO and BO) showed that DBO had a significantly lower prediction error at iteration 30 (0.04 ± 0.02, *p* = 0.048, *d*_*z*_ = 0.555). Consistent with the walking speed results, the difference between DBO and BO at the final validation iteration was no longer significant (0.03 ± 0.02 m/s, *p* = 0.159) but still in the direction of an improvement in DBO relative to BO.

Group-averaged standardized prediction error (Fig. 6 bottom) showed different trends between DBO and BO. For DBO, the standardized prediction error at each validation iteration was 10.78 ± 3.02, 6.29 ± 2.30, 4.74 ± 2.52, and 1.49 ± 0.51 for iterations 10-40, indicating a decreasing trend over iteration. In contrast, BO showed 16.79 ± 9.86, 5.27 ± 2.51, 17.88 ± 11.28, and 9.93 ± 6.07 for the same iterations, reflecting more variable changes. Paired comparisons at each iteration revealed no significant differences between the two optimizers.

## IV. Discussion

The main goal of this experiment was to implement dynamic Bayesian Optimization in a human-in-the-loop optimization problem within the context of robot-assisted gait training aimed at increasing walking speed. Sixteen healthy participants were exposed to a hip torque pulse generated by a lower extremity exoskeleton while walking on a treadmill under user-driven velocity control. The control parameters governing the hip torque profile were selected in real-time using either DBO or BO. Experimental results showed that both DBO and BO successfully increased walking speed compared to baseline (Fig. 3), with significant differences observed in paired comparisons between baseline and all validation iterations for both optimizers. As indicated by the sham torque condition in Fig. 3, this increase in speed was not primarily due to participant adaptation to a random torque condition applied by the robot, but rather driven by the optimizers identifying subject-specific optimal assistive conditions that led to increased walking speed. When comparing performance between DBO and BO, walking speed was significantly different at one of the validation iterations, as supported by the the linear mixed-model analysis, which showed a significant effect of iteration number and interaction of optimizer type and iteration number. Similarly, the prediction error was significantly different between DBO and BO at the same validation iteration,consistent with the linear mixed-model results showing interaction of optimizer type and iteration number was a significant main effect. In general, walking speed and prediction error data showed trends highlighting a general superiority of DBO relative to BO for this HILO problem, in terms of efficacy, modeling accuracy, and personalization. In terms of efficacy, DBO induced significant changes in walking speed relative to baseline that exceeded those induced by BO (Fig. 3, bottom) in three out of four validation iterations. For modeling accuracy, DBO outperformed BO in later validation iterations (Fig. 6) and improved its prediction error during cross-validation in later iterations as opposed to early iterations (Fig. 5). In terms of personalization, DBO induced changes in walking speed that were significantly greater than those induced by previously identified assistive solutions, while this was not the case of BO (Fig. 2).

The main goal of the experiment was to find individualized solutions for each participant. The general solution to increase walking speed, i.e., assisting swing or stance motion, based on findings from a previous pilot test [15], was applied during the first two iterations during the experiment. However, as shown in Fig. 2, this solution did not always have a positive effect, and in some cases, it resulted in a decrease in speed for several participants.

Therefore, the normalized difference of input parameters tested during the validation session was assessed by comparing them with the input values from iterations 1 and 2, as follows:

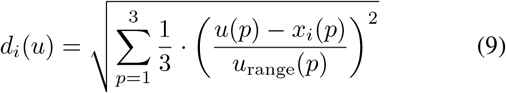

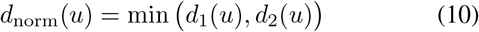

where *x*_1_ and *x*_2_ correspond to the input sets applied at iterations 1 and 2, respectively (*x*_1_ = (−9, 75, 30), *x*_2_ = (9, 50, 40)). *d*_*i*_(*u*) is a metric normalized between 0 and 1, where 0 indicates that the tested solution is identical as the one tested at the group level, while 1 indicates that the tested solution maximally differs from the one tested at the group level. *u*_*range*_ represents the range of each input parameter *T, P*, and *D* (*u*_*range*_ = (18, 100, 30)). As shown in Table I, the input parameters tested during the validation iterations showed substantial differences compared to those tested at iterations 1 and 2. This suggests that the optimizer was estimating an individualized response model, which was then tested during the validation iterations.

**TABLE I.** Normalized difference between input parameters and inputs applied at iterations 1 and 2.

| Iteration | 10 | 20 | 30 | 40 |
| --- | --- | --- | --- | --- |
| DBO | $0.26 \pm 0.05$ | $0.26 \pm 0.05$ | $0.23 \pm 0.03$ | $0.23 \pm 0.04$ |
| BO | $0.26 \pm 0.03$ | $0.36 \pm 0.05$ | $0.28 \pm 0.04$ | $0.28 \pm 0.03$ |

The internal model that estimated participant response played a crucial role in the performance of optimizer, since the performance difference between DBO and BO reflects their differing ability to account for the relevance of past observations to current conditions, which is closely tied to the model used by each optimizer to define an optimal input at each iteration during training. An example describing the training history for a selected participant is shown in Fig. 7 and 8. Analysis of these two figures (Fig. 7 and 8) suggests that DBO predicts the human-robot system response more realistically, as timing (*P*) plays a crucial role in delivering either assistive or resistive torque pulses to the participant. In contrast, the BO model shows no effect of timing on the system response. As shown in Fig. 8, the BO estimates of participant responses remained mostly constant across different sets of input parameters, resulting in large prediction and standard prediction errors, as shown in Fig. 6, as it is unable to form a sufficiently predictive model of the human response, and rather attributing the mismatch between expected and measured response to unstructured variability. An overview of model estimation for all sessions and all participant is provided in the video uploaded as supplementary materials.

**Fig. 7.**
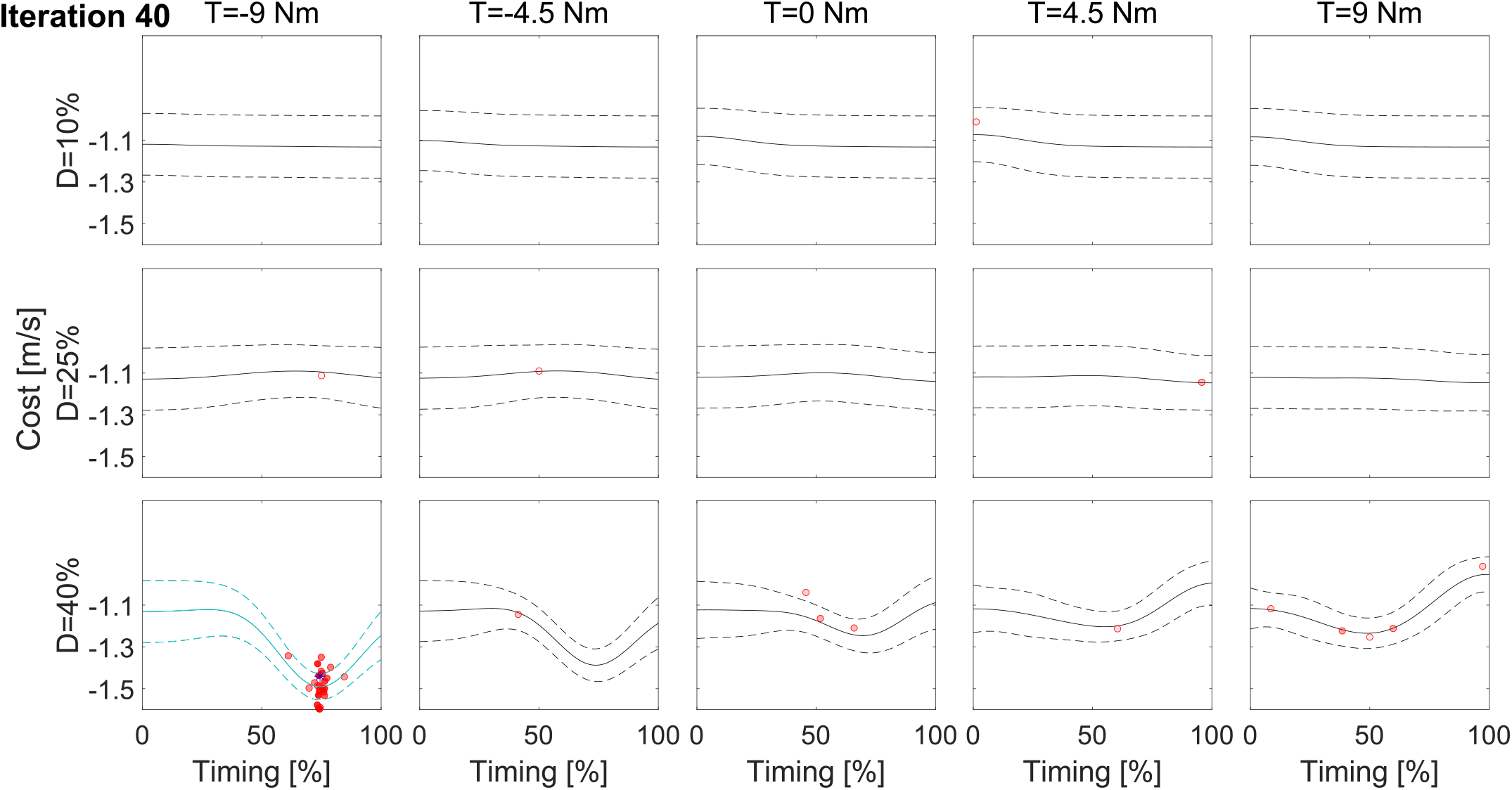
Gaussian process models for participant 2, estimating the response at iteration 40 during training with DBO as the optimizer. The models were constructed using the response history prior to the validation sessions at iteration 40. The solid line represents the estimated mean cost function, with dashed lines indicating one standard deviation. Each subplot displays the estimated Gaussian model, with the torque amplitude (*T*) indicated in the title, duration (*D*) on the y-axis, and timing (*P*) on the x-axis. The cyan line represents the Gaussian process model evaluated at the torque amplitude (*T*) and duration (*D*) tested in the 40^th^ iteration. The model is shown within a window corresponding to the closest values of *T* and *D*, as indicated in the title and y-axis. Dots represent past responses, with greater transparency indicating earlier iterations, and are displayed within the same window defined by the closest values of *T* and *D*. Asterisks indicate the actual response at the 40^th^ iteration.

**Fig. 8.**
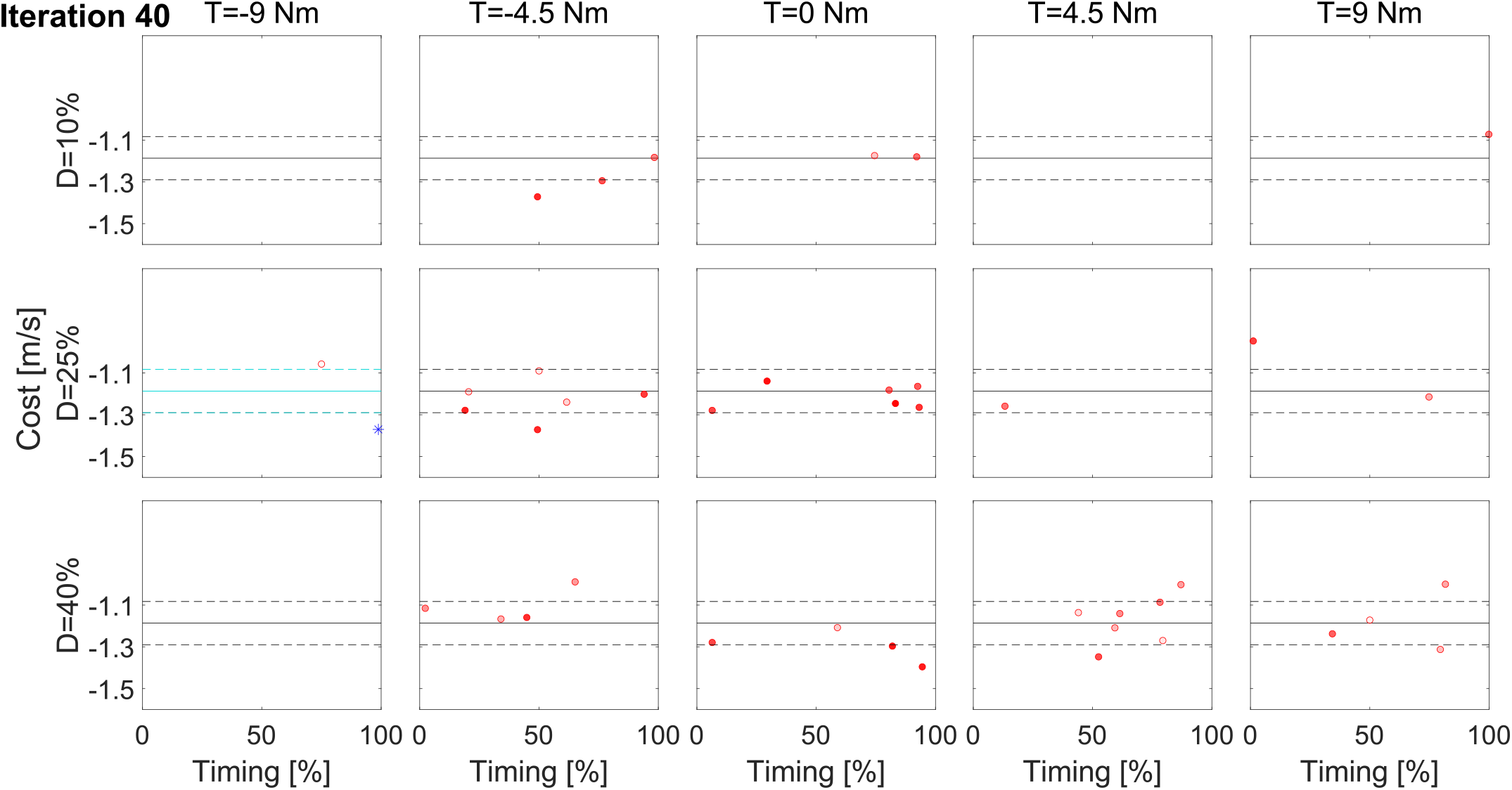
Gaussian process models for participant 2, estimating the response at iteration 40 during training with BO as the optimizer.

The non-stationarity of the human-robot system was assumed to arise naturally from the experimental setup. At each iteration, participants experienced a torque pulse for 40 strides, followed by a no-torque condition for 20 strides. However, some participants required more strides to fully wash out the effects of the torque pulses they had previously experienced. As a result, the next iteration, which involved a new torque pulse, started with a different walking speed compared to the previous iteration. Moreover, with training, participants would be able to take advantage of the exoskeleton assistance in different ways, leading to their response not always being consistent across iterations. This overall variability in responses made it challenging for the optimizer to accurately predict responses. As shown in Fig. 8, BO was unable to effectively weigh past observations in relation to current ones. Therefore, when similar inputs were applied, BO treated different responses (caused by varying initial speeds and participant response variability) the same way. This led to the GP model failing to predict the control parameters effectively.

Several studies have investigated the use of lower-limb exoskeletons in relation to changes in user walking speed. However, these studies often did not directly target walking speed as the primary outcome. Instead, they focused on adapting exoskeleton control strategies in response to changes in walking speed that had already occurred [25], or implemented predefined control strategies in post-stroke individuals and subsequently evaluated their effects on walking speed [26], [27]. Previous implementation of HILO with an ankle exoskeleton successfully identified individualized control solutions that increased walking speed [28], with the resulting changes in walking speed generally exceeding the one demonstrated in this study (0.52 m/s; 0.30 m/s in this study). However, participants were required to complete two continuous 72-minute walking sessions to identify the optimal parameters, which may limit the feasibility of this approach for rehabilitation applications due to potential fatigue and endurance constraints. The results of the present study demonstrate the feasibility and potential of implementing HILO with DBO in a hip exoskeleton, particularly due to its capability to accommodate the non-stationary characteristics of human–robot interactions. Furthermore, the proposed method exhibited superior performance compared with conventional BO, as demonstrated through multiple validation iterations that provide insight into the optimizer’s behavior during and after the training process. The findings suggest that this approach may provide a more practical and efficient framework for identifying individualized assistance strategies while reducing the burden associated with prolonged optimization procedures.

## V. Conclusion

In this experiment, we aimed to implement DBO and BO within the HILO framework for robot-assisted gait training to improve walking speed. Both DBO and BO successfully induced increases in walking speed compared to baseline, with significant improvements observed in paired comparisons between baseline and all validation iterations for both optimizer types. When comparing the performance of DBO and BO, significant differences were found only at one of the validation iterations for both walking speed and prediction error, with linear mixed model analysis results showing a significant effect of interaction between optimizer type and iteration number on both the walking speed and prediction error. Overall, while both DBO and BO were able to increase walking speed, DBO showed more promise in terms of prediction accuracy and adaptability, particularly in the context of human-in-the-loop optimization where individual variability and system non-stationarity are critical factors.

There are several limitations to this study. First, the total number of iterations was limited to 40, which may not have been sufficient to fully observe and quantify differences in performance between DBO and BO. Since the optimizer targeted walking speed, which requires several strides for the participant to achieve a stable response under each torque intervention, it was necessary to increase the number of strides per iteration. This, in turn, limited the total number of iterations in order to avoid participant fatigue. Also, the maximum treadmill speed was restricted to 1.6 m/s for safety reasons, which posed a challenge for the optimizer, particularly when participants were walking close to this maximum speed regardless of the input parameters being tested. This was the case only for some of the participants in our study (Participant 1 with DBO, and Participant 7 with BO). These near-constant responses may have led the optimizer to underestimate the effect of the torque pulses on the participants. Furthermore, the analysis pursued to quantify the effect of either control algorithm does not account for the specific input parameters tested in each iteration, but instead captures only general trends with respect to iteration number. This limitation does not allow to understand how each input value evaluated by the optimizer influenced the outcome.

## Notes

This work is supported in part by NIH-R01HD111071 and in part by American Heart Association under Grant TPA – 947225.

### Competing Interest Statement

The authors have declared no competing interest.

